## Supplemental Material for "CONGA: Copy number variation genotyping in ancient genomes and low-coverage sequencing data"

### 1. Command Lines

#### 1.1. Simulations

##### 1.1.1. Varsim [1]

```
./varsim.py --vc_in_vcf tests/quickstart_test/21_5_10Mb.vcf.gz --sv_insert_seq
insert_seq.txt --sv_dgv GRCh37_hg19_supportingvariants_2013-07-23.txt
--reference $REFERENCE_GENOME --id varsimu --sv_num_del 2000
--sv_num_dup 2000 --sv_percent_novel 0.01 --sv_min_length_lim 1000
--sv_max_length_lim 10000 --total_coverage 10 --java_max_mem 50g
--simulator_executable /home/asoylev/art_bin_GreatSmokyMountains/art_illumina
```

##### 1.1.2. Gargammel [2]

```
perl gargammel.pl -c $COVERAGE --comp 0,0,1 -damage 0.024,0.36,0.0097,0.55 -f
src/sizefreq.size.gz -s /usr/local/sw/gargammel/src/sizedist.size.gz -ss HS25 -o
$OUTPUT
```

##### 1.1.3 Adapter Removal and Merge [3]

```
AdapterRemoval --file1 $LEFT_PAIR --file2 $RIGHT_PAIR --qualitybase 33 --gzip
--qualitymax 60 --trimns --collapse --minalignmentlength 11 --basename $FILENAME
--settings $FILENAME.settings

cat $FILENAME.collapsed.gz $FILENAME.collapsed.truncated.gz
$FILENAME.pair1.truncated.gz
$FILENAME.pair2.truncated.gz>$FILENAME.all.fastq.gz
```

#### 1.2. Alignment

**Alignment to the reference genome [4]:**

```
bwa aln -l 16500 -n 0.01 -o 2 -t ${cores} ${REFERENCE_GENOME}
${mergedname} \
| $bwa samse ${REFERENCE_GENOME} - ${mergedname} \
| samtools view -F 4 -h -Su - \
| samtools sort - -o ${alndir}/mapped/${filebase}.${refbase}.bam
```

**Filter out PCR duplicates (FilterUniqueSAMCons_rand.py)** [5]:

```
samtools view -F 4 -h ${alndir}/mapped/${filebase}.${refbase}.bam \
| python ${scriptdir}/FilterUniqueSAMCons_rand.py | samtools view -h -Su - \
> ${alndir}/mapped/${filebase}.${refbase}.cons.bam
```

**Filter out high mismatch and short reads (percidentity_threshold.py):**

```
samtools calmd ${alndir}/mapped/${filebase}.${refbase}.cons.bam
${REFERENCE_GENOME} \
| python ${scriptdir}/percidentity_threshold.py 0.9 35 ${TMPDIR}/short.txt \
| samtools view -bS - > ${alndir}/${filebase}.${refbase}.cons.90perc.bam
```

#### 1.3. CNV Discovery and Genotyping

**CONGA:**

```
conga -i $BAM_FILE --sonic $SONIC_FILE --ref $REFERENCE_GENOME
--dels $DELS_INPUT --dups $DUPS_INPUT --mappability
$MAPPABILITY_FILE -o $OUTPUT
```

**CNVNator** [6] **:**

```
cnvnator -root file.root -tree $BAM_FILE -chrom 1 2 3 4 5 6 7 8 9 10 11 12 13 14 15 16 17 18 19 20 21 22 X Y MT

cnvnator -root file.root -his 1000 -d /ref/

cnvnator -root file.root -stat 1000

cnvnator -root file.root -partition 1000

cnvnator -root file.root -call 1000 >$OUTPUT
```

* Note that we used window of 100 for BAM files with small variations and 1000 in all the other BAMs.

**FREEC** [7] **:**

```
freec -conf data/config_WGS_human.txt
```

* Default configurations were used in the config file of FREEC

**GenomeSTRiP** [8]**:**

```
java -cp $SV_CLASSPATH ${javaMaxMemory} \
org.broadinstitute.gatk.queue.QCommandLine \
-S ${SV_DIR}/qscript/SVPreprocess.q \
-S ${SV_DIR}/qscript/SVQScript.q \
-configFile ${SV_DIR}/conf/genstrip_parameters.txt \
-tempDir tmp \
-gatk ${SV_DIR}/lib/gatk/GenomeAnalysisTK.jar \
-cp ${SV_CLASSPATH} \
-jobLogDir metadata/logs \
-R $REFERENCE_GENOME \
-md metadata \
-ploidyMapFile ${ploidy} \
-I ${bamFileList} \
-genomeMaskFile  $REFERENCE_GENOME_MASK36 \
-copyNumberMaskFile  $REFERENCE_GENOME_CN2_MASK \
-bamFilesAreDisjoint true \
-computeGCProfiles true \
-deleteIntermediateDirs true \
```

```
-jobRunner ParallelShell \
-run \

java -cp ${SV_CLASSPATH} -Xmx4g \
org.broadinstitute.gatk.queue.QCommandLine \
-S ${SV_DIR}/qscript/SVGenotyper2.q \
-S ${SV_DIR}/qscript/SVQScript.q \
-gatk ${SV_DIR}/lib/gatk/GenomeAnalysisTK.jar \
-cp ${SV_CLASSPATH} \
-configFile ${SV_DIR}/conf/genstrip_parameters.txt \
-jobLogDir ${runDir}/logs \
-tempDir ${SV_TMPDIR} \
-R ${referenceFile} \
-md ${mdPath} \
-genomeMaskFile $REFERENCE_GENOME_MASK36 \
-genderMapFile ${SV_DIR}/input/sample_gender.report.txt \
-I ${bamFileList} \
-parallelRecords 1000 \
-runDirectory ${runDir} \
-vcf ${vcfFile} \
-skipAnnotator CNQuality \
-O /${runDir}/output_file.genotypes.vcf \
-jobRunner ParallelShell \
-gatkJobRunner ParallelShell \
-run
```

* We used default mask files for human genome grch37 with k-mer size 36.

#### mrCaNaVaR

**Alignment using mrsFAST** [9]:

```
mrsfast --search $MASKED_REFERENCE_GENOME --seq $FASTQ_FILE
--seqcomp --threads 24 -o sim.sam --disable-nohits --crop 36
```

**Variant Calling using mrCaNaVaR** [10, 11]**:**

```
mrcanavar --read -conf human_g1k_v37.cnvr -samlist $SAM_FILE -depth
$DEPTH_FILE_NAME

mrcanavar --call human_g1k_v37.cnvr -depth $DEPTH_FILE_NAME -o
$OUTPUT_FILE
```

#### 1.4. BEDTools [12]

**TRUE_PREDICTION** = ~/bedtools2/bin/intersectBed -a $TRUE_CALLS -b del.bed -f 0.5 -wa -r|grep -v X|grep -v Y|grep -v GL|grep -v MT|sort -u|wc

**FALSE_PREDICTION** = ~/bedtools2/bin/intersectBed -a del.bed -b $TRUE_CALLS -f 0.5 -r -wa -v|grep -v GL|grep -v X|grep -v Y|grep -v MT|sort -u|wc

**FDR** = FALSE_PREDICTION / (TRUE_PREDICTION + FALSE_PREDICTION)

**RECALL** = TRUE_PREDICTION / (TRUE_PREDICTION + FALSE_PREDICTION)

**PRECISION** = TRUE_PREDICTION / TRUE_CALLS

**F-SCORE** = (2 * PRECISION * RECALL) / (PRECISION + RECALL)

#### 1.5 Depth of coverage calculation [13]

samtools view -b -q 30 -F 4 XXX.bam | genomeCoverageBed -ibam - -g human_g1k_v37.fasta >coverage.cov

grep genome coverage.cov | awk '{NUM+=$2*$3; DEN+=$3} END {print NUM/DEN}'

#### 1.6 Down-sampling

java -jar ~/picard.jar DownsampleSam I=XXX.bam O=XXX_0004.bam P=0.004

* In our experiments, we used “P” values of 0.7, 0.3, 0.1, 0.05, 0.03, 0.01, 0.04, 0.03 for Yamnaya, 0.7, 0.3, 0.1, 0.05, 0.03, 0.01, 0.04 for Saqqaq and 0.7, 0.3, 0.1, 0.05, 0.03 for Mota genomes.

### 2. Supplemental Figures

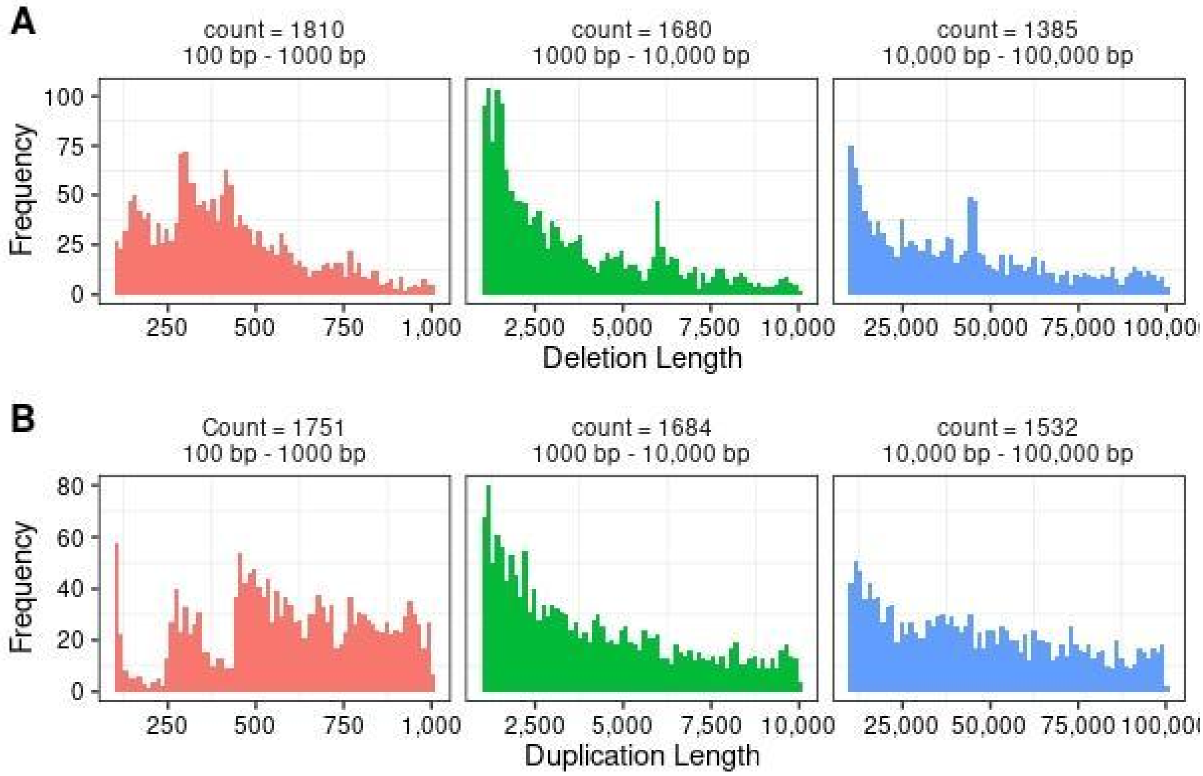


**Supplemental Figure S1**. Length distributions and numbers of deletions (A) and duplications (B) inserted into three simulated genomes. The total number of CNVs inserted into a genome ("counts") is shown at the top of each graph. We used Varsim to insert these CNVs into each genome, yielding three genomes in total (for short, medium and large CNVs).

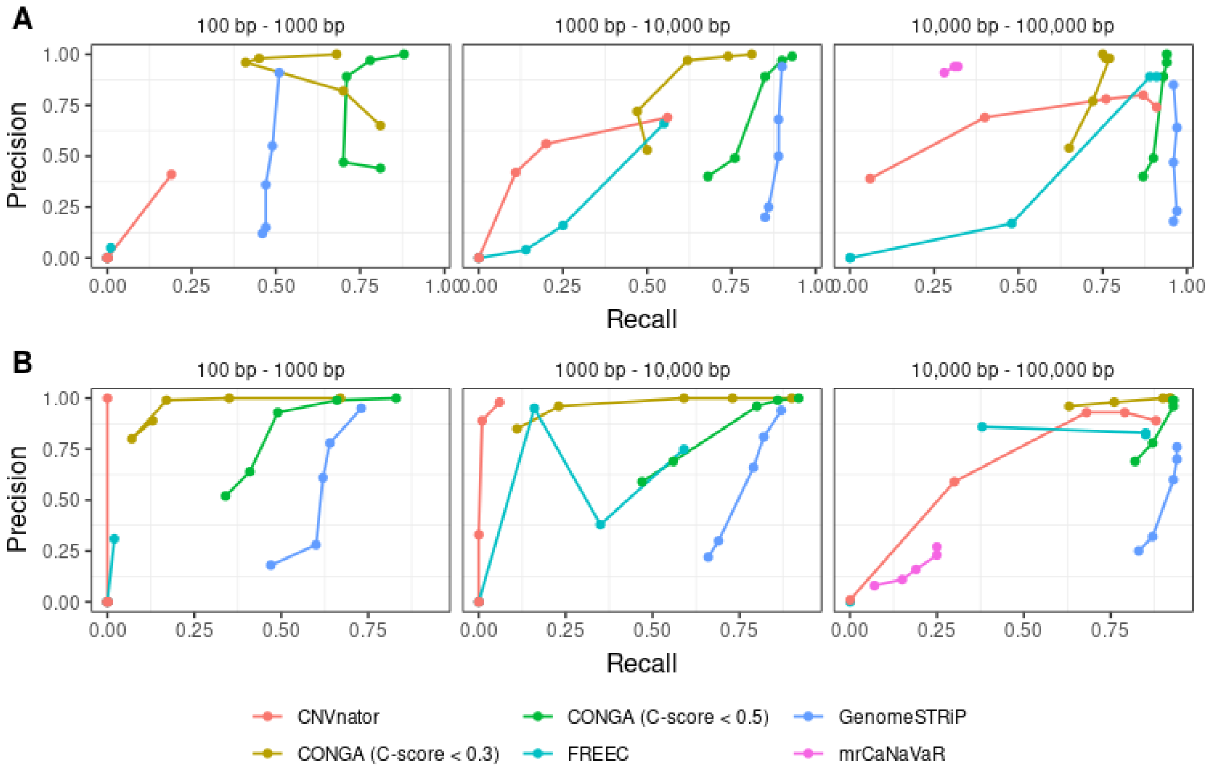


**Supplemental Figure S2**. Precision-Recall curves for deletion (A) and duplication (B) predictions of CONGA, GenomeSTRiP, FREEC, and CNVnator using coverages of 0.05x, 0.1x, 0.5x, 1x and 5x. mrCaNaVaR was used only in the analysis of large variants.

**A**

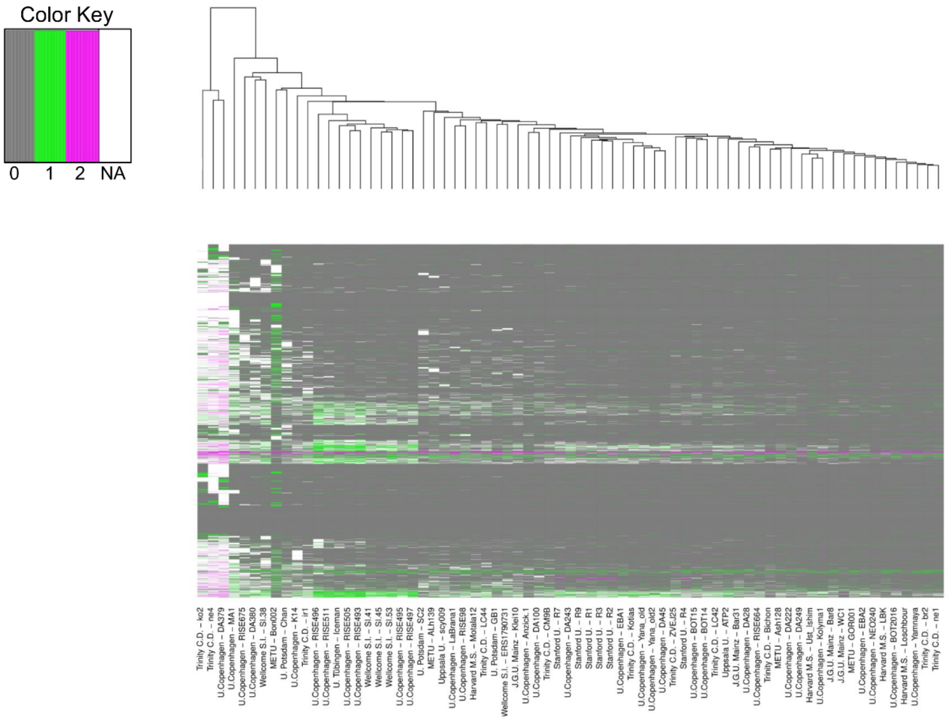


**B**

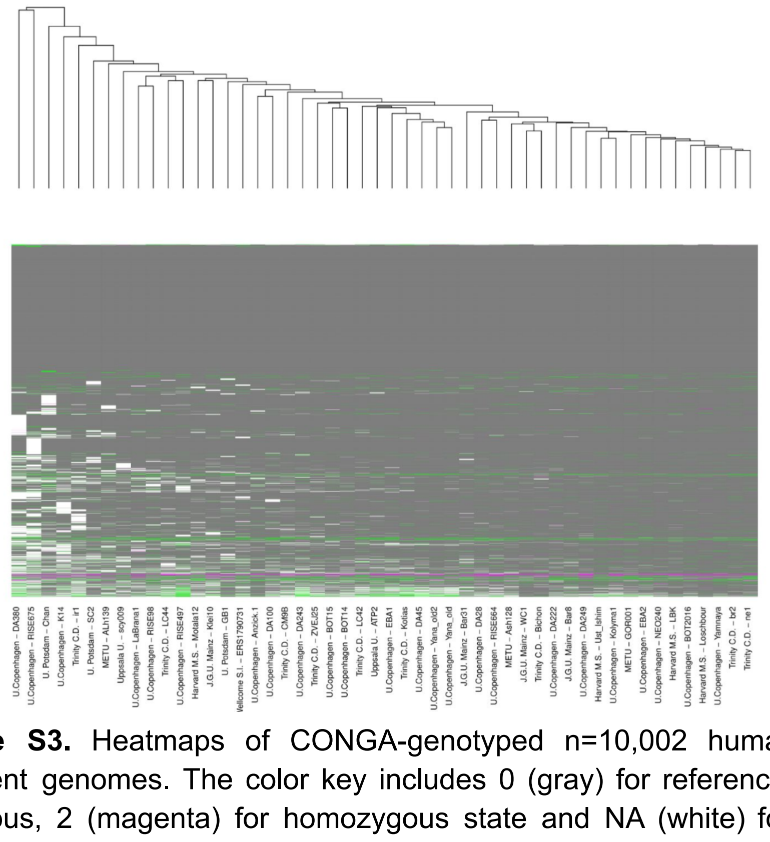

**Supplemental Figure S3.** Heatmaps of CONGA-genotyped n=10,002 human-derived deletions across ancient genomes. The color key includes 0 (gray) for reference allele, 1 (green) for heterozygous, 2 (magenta) for homozygous state and NA (white) for missing value. (A) Heatmap of deletions per genome on the raw dataset (n=71 genomes). (B) Heatmap of deletions per genome on the refined dataset (with n=50 genomes after removing divergent genomes).

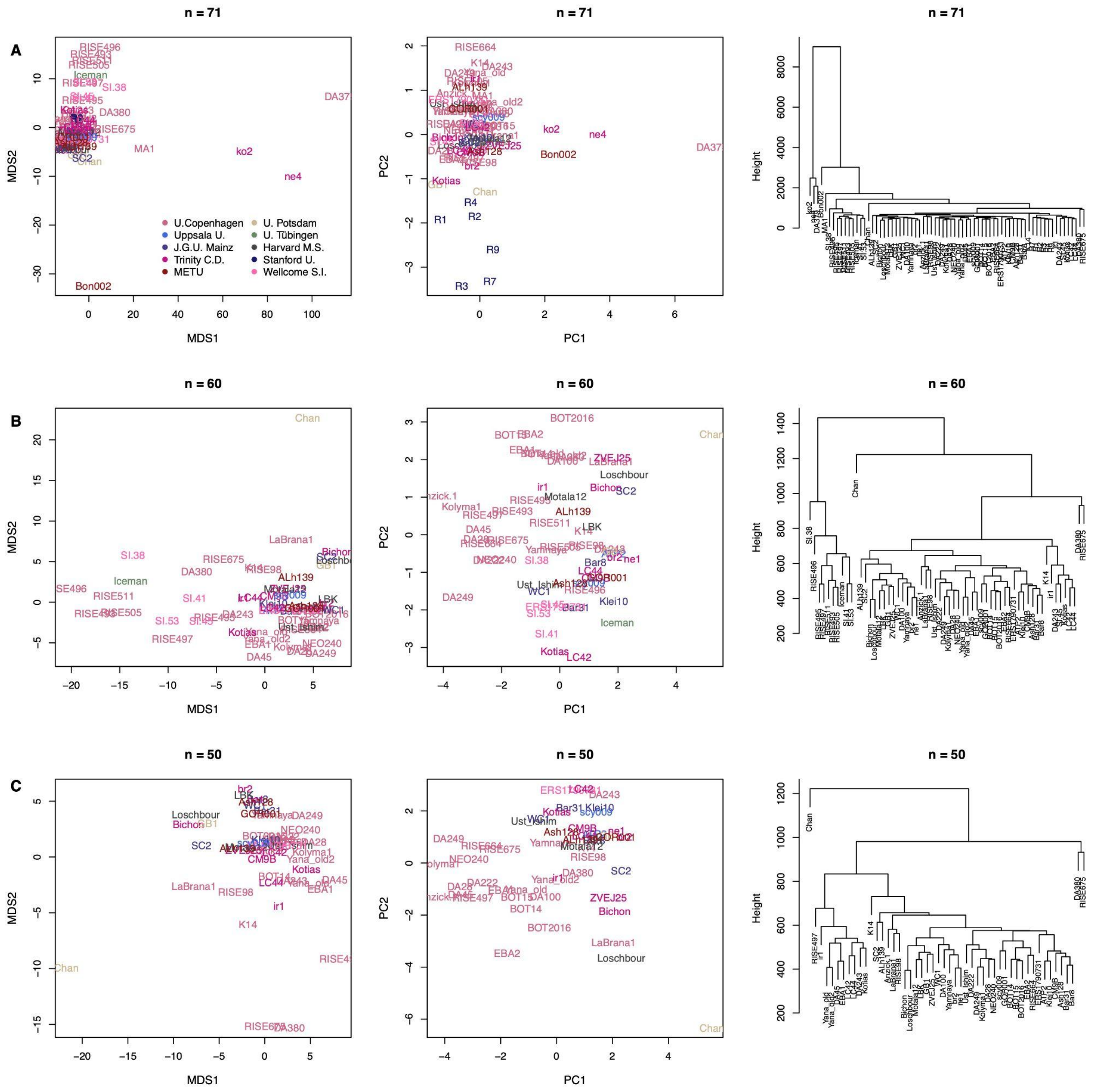


**Supplemental Figure S4**. Multivariate analysis of deletion frequencies reveal outlier genomes. Left panels: Multidimensional scaling plots (MDS) calculated with k=2 using the R "cmdscale" function on a Euclidean distance matrix of deletion frequencies. Middle panels: Principal component analysis plots (PCA) summarizing deletion frequencies after removing any NAs. Right panels: Hierarchical clustering trees summarizing Manhattan distance matrices, calculated using the R "dist" and "hclust" functions. The color codes indicate the laboratory-of-origin of each genome, shown in the legend of the top right panel. (A) Results based on the full dataset with 10,002 human-derived deletions (n=8,780 genotyped in any state in at least one genome) and n = 71 genomes. In the PCA we use $n_D$ = 580 deletions after removing loci with at least one missing value. (B) Results based on n = 60 genomes after removing 11 outlier genomes (and $n_D$ = 3,460 deletions in the PCA). (C) Results based on n = 50 genomes after removing 21 outlier genomes (and $n_D$ = 3,472 deletions in the PCA). We note that the MDS here differs from that shown in Fig. 4, in that the latter is calculated using outgroup-$f_3$ statistics.

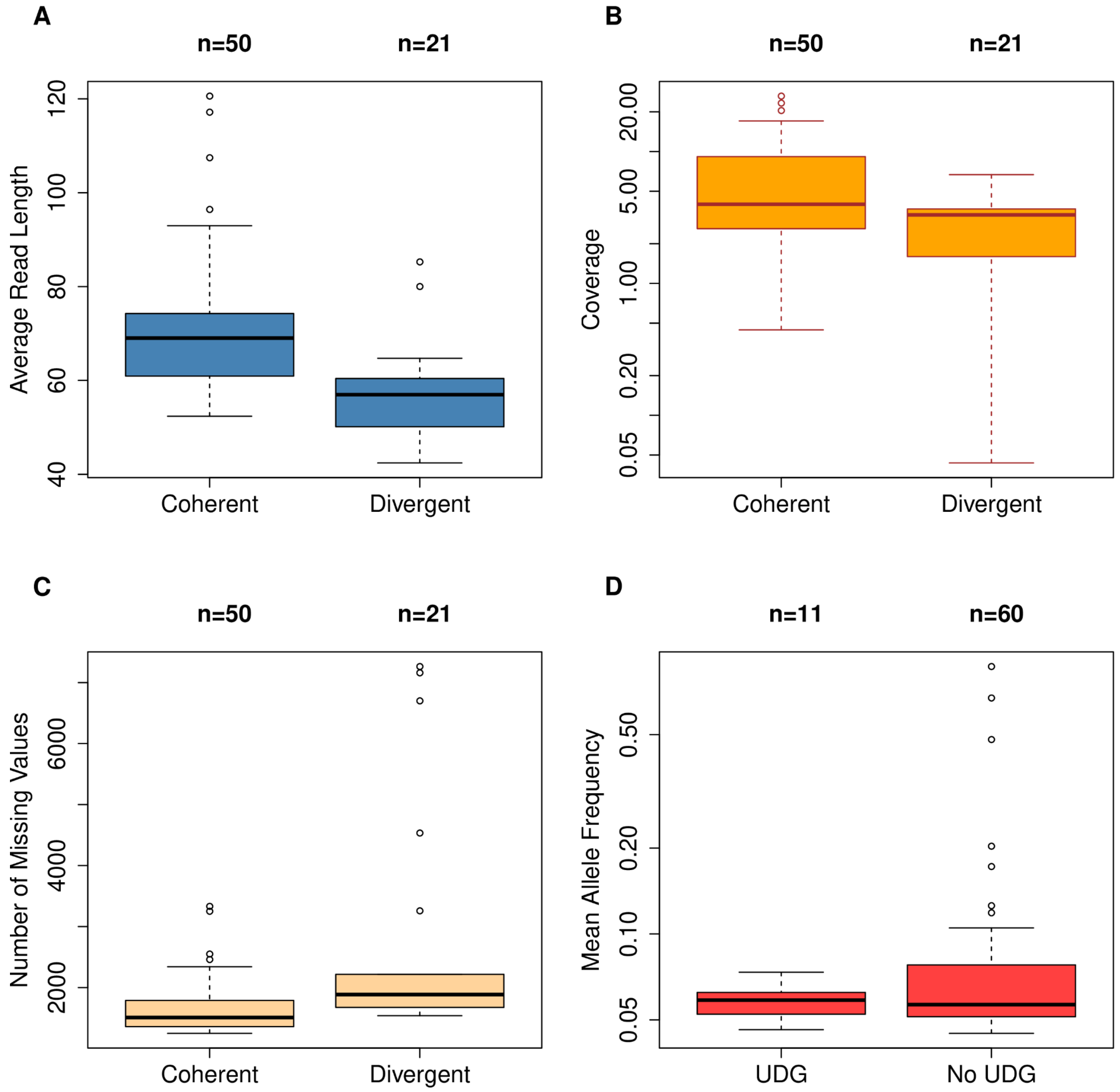


**Supplemental Figure S5**. Technical comparisons of the divergent (n=21) and coherent (n=50) genome sets, defined based on their deletion profiles (Supplemental Figure S3 and Supplemental Figure S4). (A) Boxplots of the average read length per genome (Wilcoxon-rank sum test, *P*<0.001). (B) Boxplots of mean depths of coverage (Wilcoxon rank sum test, *P*=0.014). (C) Boxplots of the number of missing values per genome (Wilcoxon-rank sum test, *P*<0.001). (D) Boxplots of the mean deletion allele frequencies per UDG-treated and not UDG-treated genomes. We observe no significant difference between the distributions (Wilcoxon-rank sum test, *P*=0.58).

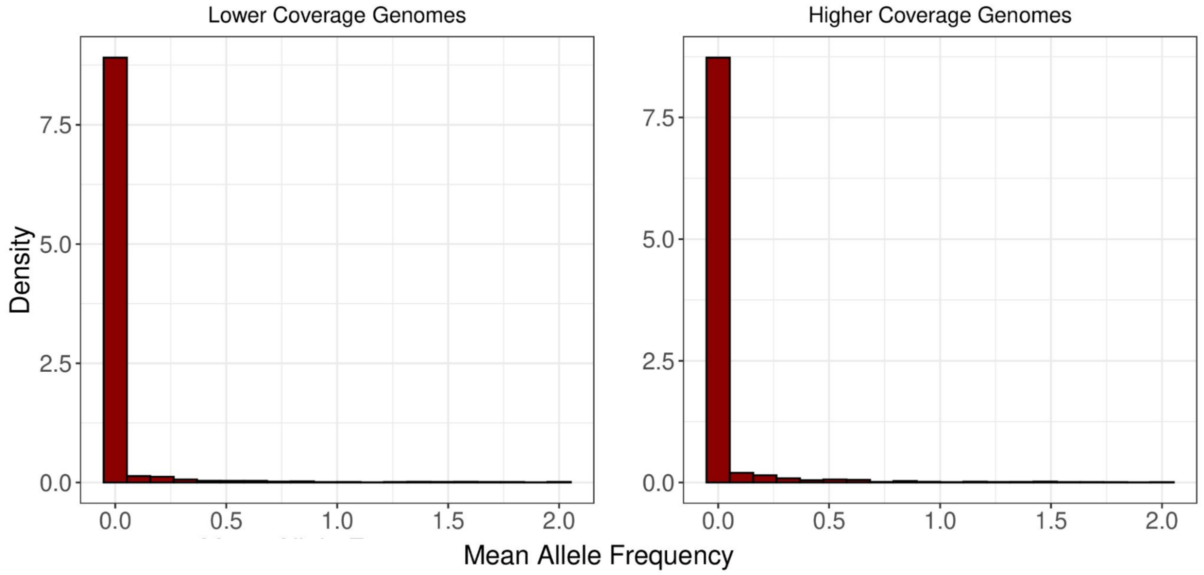


**Supplemental Figure S6.** Site-frequency spectra of deletions genotyped in low and high coverage genomes. The left panel represents the SFS of n=25 below-median coverage genomes and the right panel shows the SFS of n=25 above-median coverage genomes. The median coverage value was 3.98x. We found no significant difference between the two SFS distributions (Kolmogorov-Smirnov test p=0.27).

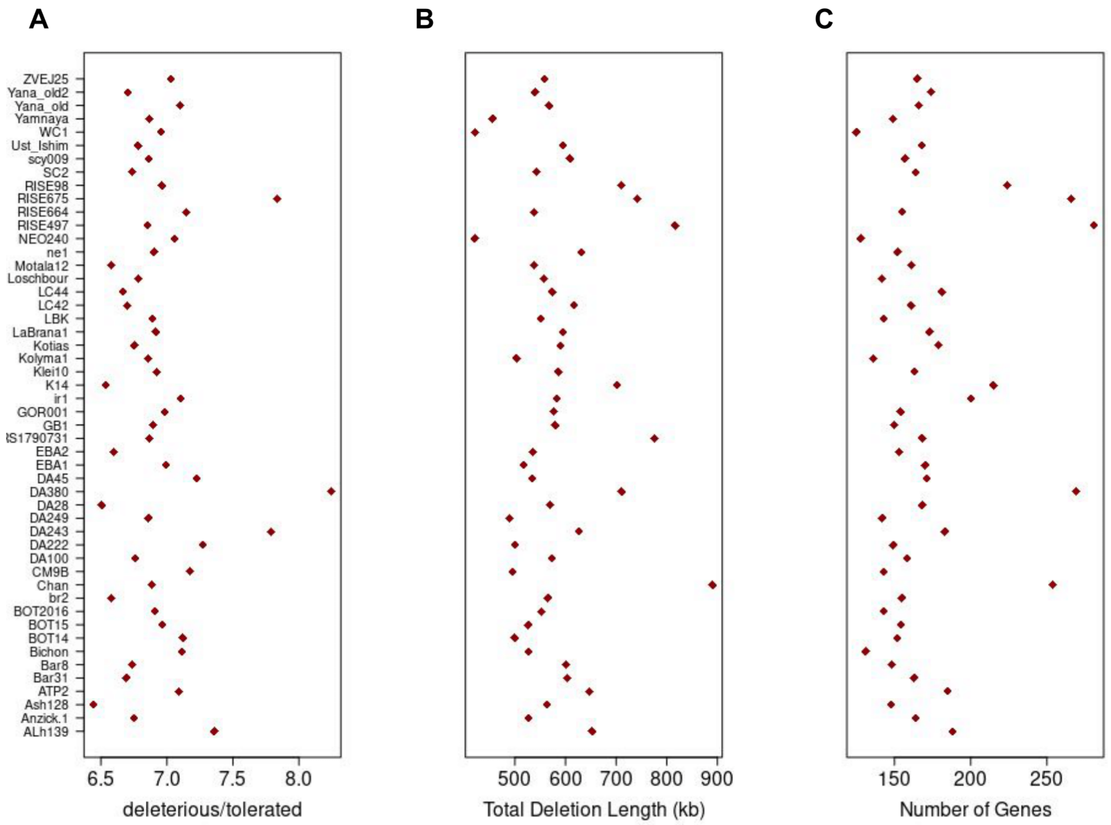


**Supplemental Figure S7.** Deleterious load estimates among 50 ancient genomes. In all three panels, the x-axis represents a deleterious load-related statistic and the y-axis shows the ancient individuals. (A) Deleterious load based on SIFT-estimated SNP effects per individual. The x-axis represents the number of “deleterious” SNPs over the number of “tolerated” SNPs. (B) CONGA-estimated total deletion length in kb per individual, using the Final CNV call-set. (C) The number of genes that overlap with CONGA-estimated deletions. In panels B and C, heterozygous and homozygous calls were counted once. In panel C, the most affected individuals in terms of the number of gene overlaps are RISE497 (Russia, 2nd millennium BCE), DA380 (Turkmenistan, 4th millennium BCE), RISE675 (Russia, 3rd millennium BCE), and Chan (Iberia, 8th millennium BCE). We observed that these individuals were around 50% more affected than the rest.

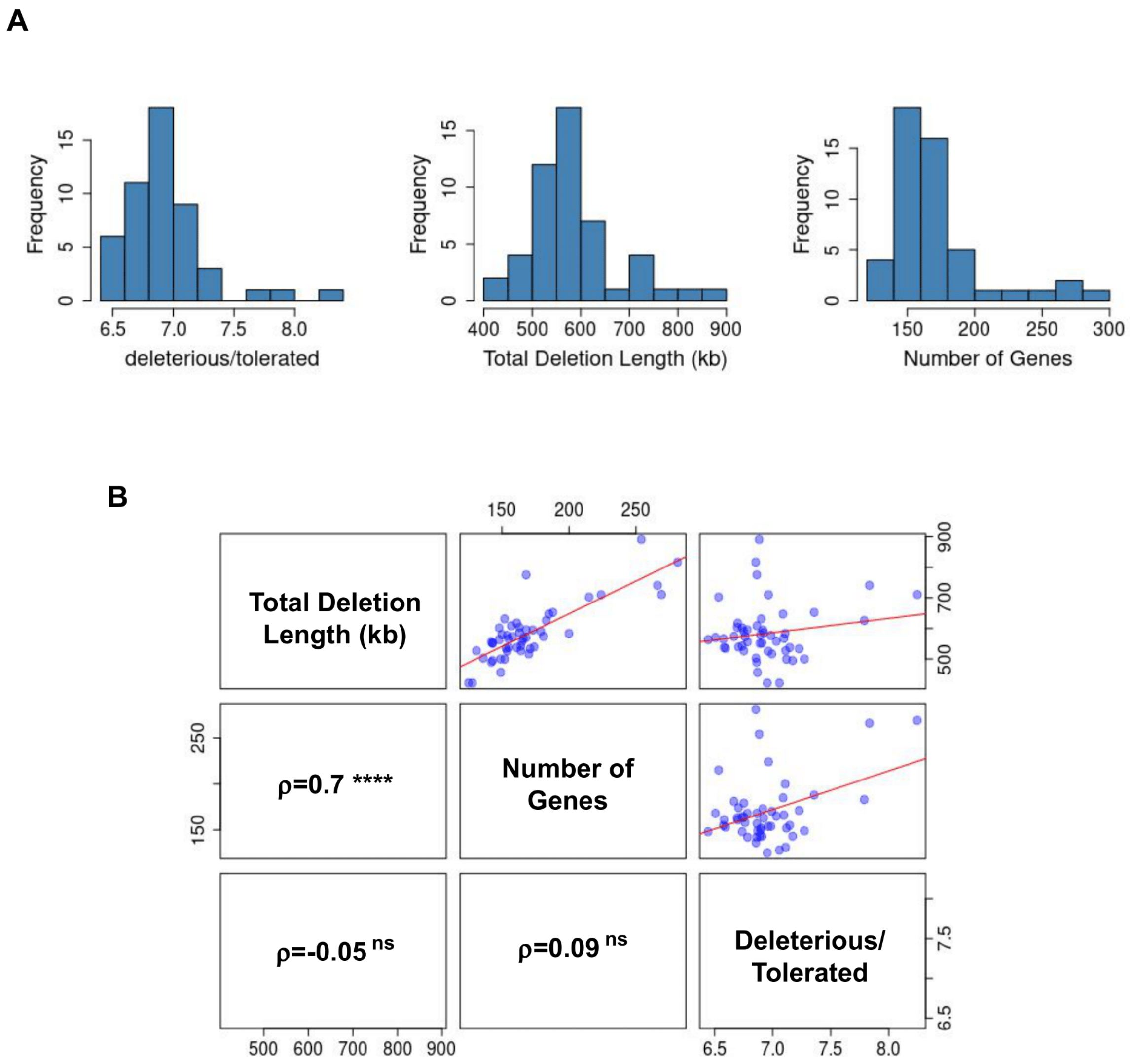


**Supplemental Figure S8.** Correlations between SNP- and deletion-based deleterious load estimates in 50 ancient genomes. (A) From left to right: histograms of the number of SIFT-predicted "deleterious" SNPs over "tolerated" SNPs per genome, CONGA-predicted total deletion length in kb per genome, and the number of genes that overlap with CONGA-predicted deletions per genome. (B) Correlations between each variable. The RHS triangle shows the scatter plots between two variables, and the LHS triangle shows the Spearman rank correlation estimates. The significance of the ρ's are also shown. **** $p$<0.0001, ns: non-significant.

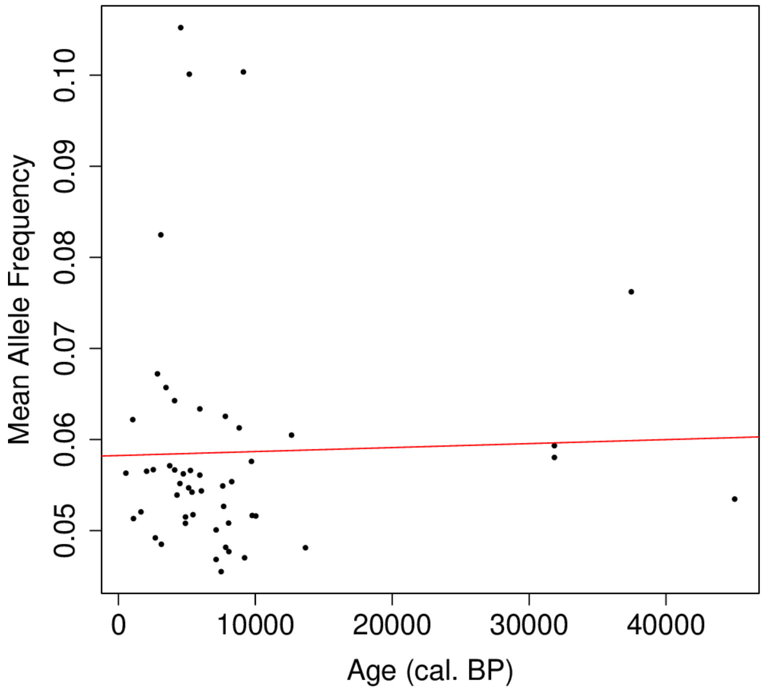


**Supplemental Figure S9.** The scatter plot of the mean allele frequencies per genome (n=50) vs. age of the genomes (calibrated years before present). The red line represents the linear model. We found that there is no significant correlation between the age of the individuals and the mean allele frequency (Spearman's rank correlation $\rho$ = -0.12, $P$ = 0.41).

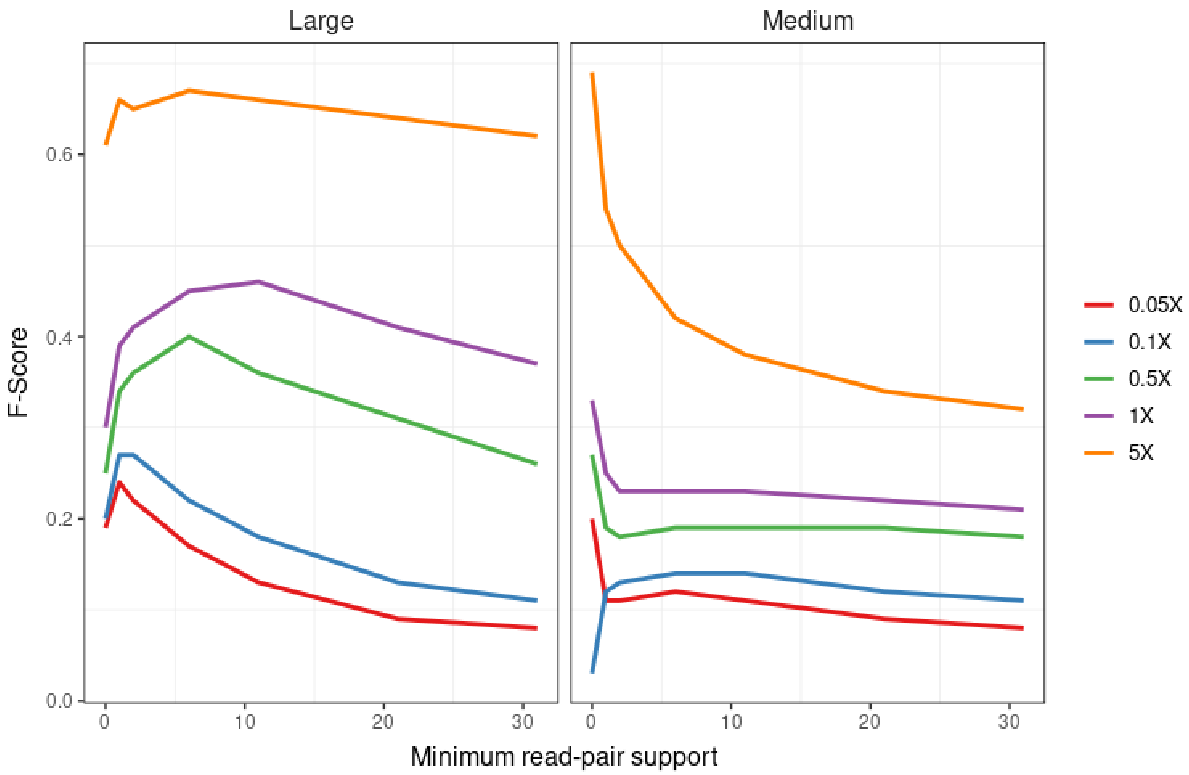


**Supplemental Figure S10.** The figure shows the effect of minimum read-pair support on the F-Score for duplications using various depths of coverage values in simulated genomes. Medium sized CNVs are between 1,000 bps to 10,000 bps and large CNVs are between 10,000 bps and 100,000 bps. Here, we used a relaxed C-score threshold of 10 in order to observe the effect of read-pair support only. The figure shows that read-pair support is effective when the coverage is above 0.5x and also when the duplication sizes are larger.

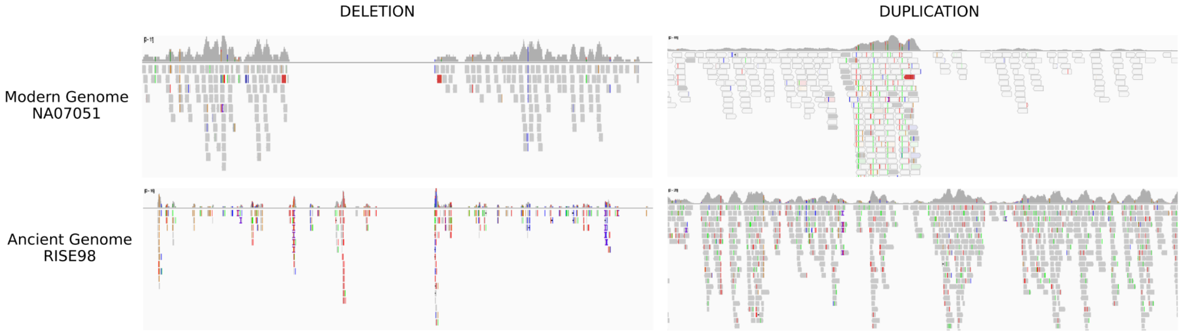


**Supplemental Figure S11**. The figure shows IGV visualization of high scoring (i.e., high likelihood) deletions (left) and duplications (right) predicted by CONGA. The events displayed in the upper panels were detected in a modern-day human genome (NA07051: an ~8 Kbp deletion within chr7:16,169,440-16,177,556 and a ~4 Kbp duplication within chr7:22,496-26,553) and those in the lower panels in an ancient genome (RISE98: an ~17 Kbp deletion within chr6:32,506,809-32,524,264 and a ~6 Kbp duplication within chr1:1,520,604-1,526,959). The candidate CNV list used for genotyping was the long read CNV dataset described in Methods. Deducing the CNVs is straightforward with the modern-day genome data, however, it is less straightforward to distinguish these variations in ancient read data, especially for duplications. Note that this is one of the sample scenarios and we emphasize that a large number of CNVs identified in ancient genomes suffer from the same issue.

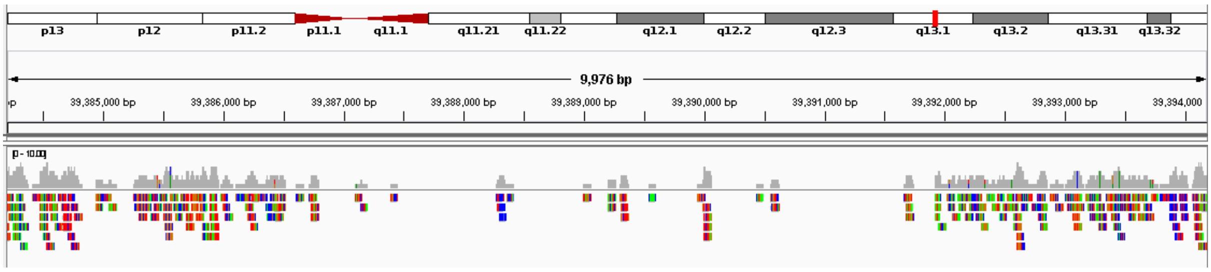


**Supplemental Figure S12**. An inserted deletion in a simulated ancient genome at 1x coverage. The event location is chr22:39,386,521-39,391,930. CONGA missed this deletion due to the poor signal.

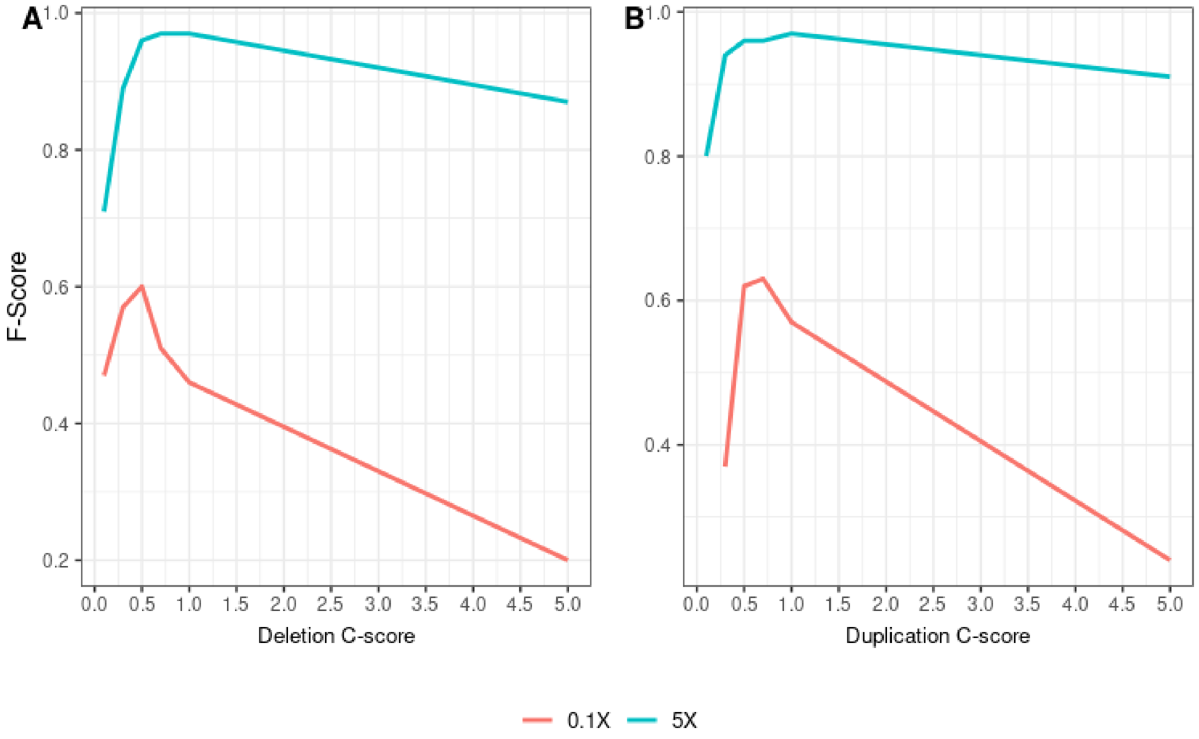


**Supplemental Figure S13**. The figure shows the effect of C-score on accuracy F-Scores of deletions (A) and duplications (B) for 0.1x and 5x depths of coverages in simulated genomes with medium sized CNVs embedded. Here, we did not use read-pair support or mappability filtering, in order to only test the effect of the C-score threshold. The C-score is calculated using read-depth information.

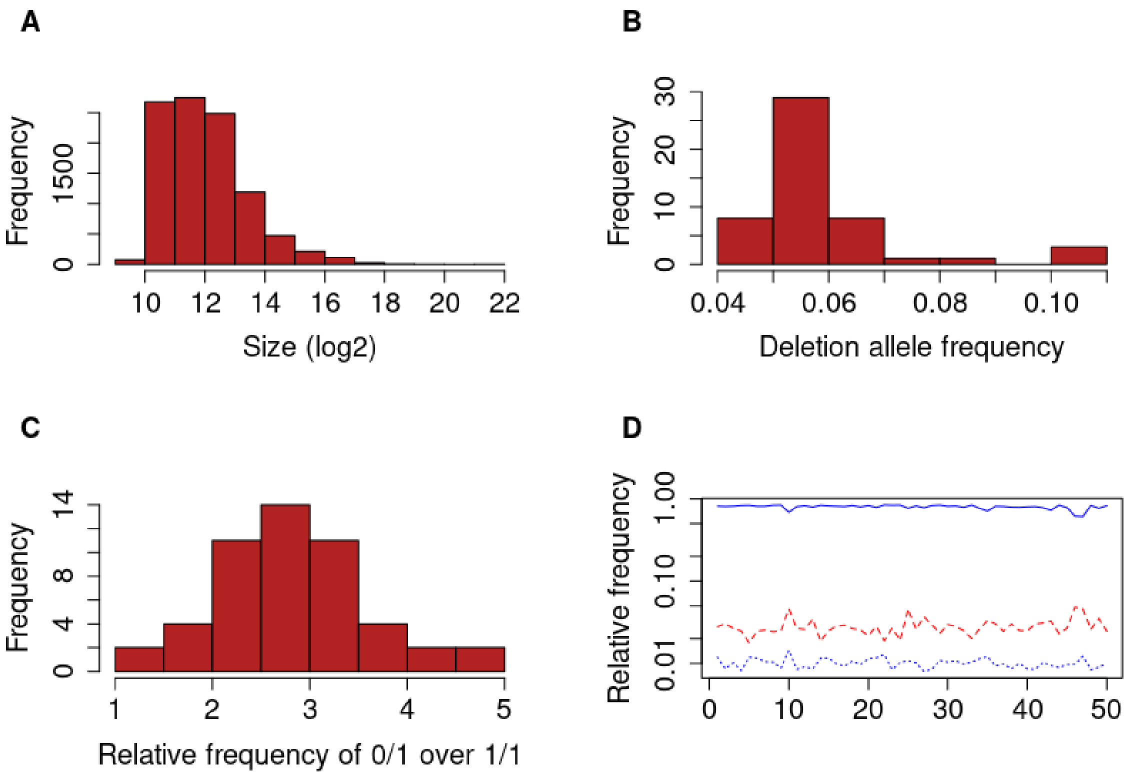


**Supplemental Figure S14.** General characteristics of deletions in the refined dataset (n=50 genomes and n=8,780 deletions, obtained after applying ancestry state filters and removing outlier genomes). (A) Size distribution of the deletions in logarithmic scale. (B) The distribution of the deletion allele frequency (i.e. the proportion of deletion alleles across the 8,780 loci per genome) among the 50 genomes. (C) The distribution of the relative frequency of observed heterozygous (0/1) deletions over homozygous (1/1) deletions observed in our dataset. (D) The plot of relative deletion frequencies called heterozygous (red lines) and homozygous (blue lines) among 8,780 deletions, for each of the n=50 ancient genomes in the refined dataset (after applying additional ancestry state filters and removing outlier genomes).

### 3. Supplemental Table Legends

**Supplemental Table S1A**. The table shows the CNV predictions of CONGA, GenomeSTRiP, FREEC, CNVnator and mrCaNaVaR on simulated genomes at depths 0.05x, 0.1x, 0.5x, 1x and 5x for deletions and duplications of multiple CNV size intervals including 100 bps - 1 Kbps (small), 1 Kbps - 10 Kbps (medium) and 10 Kbps - 100 Kbps (large). Here, “T” and “F" refer to correct and incorrect predictions respectively, “Miss” is the number of missed true events, “Recall” (TPR) is the true positive rate, and “FDR” is false discovery rate (1 - “Precision”) for each run. The F-Score is calculated as (2 * Precision * Recall) / (Precision + Recall). Note that for CONGA, we included the performance for both C-score<0.5 and C-score<0.3.

**Supplemental Table S1B**. The table shows the copy-number (homozygous or heterozygous) predictions of CONGA on simulated genomes at depths 0.05x, 0.1x, 0.5x, 1x and 5x for deletions and duplications of multiple CNV size intervals including 100 bps - 1 Kbps (small), 1 Kbps - 10 Kbps (medium) and 10 Kbps - 100 Kbps (large). Here, “T” and “F” refer to correct and incorrect predictions respectively, “Miss” is the number of missed true events, “Recall” (TPR) is the true positive rate, and “FDR” is false discovery rate (1 - “Precision”) for each run. The F-Score is calculated as (2 * Precision * Recall) / (Precision + Recall). Note that for CONGA, we included the performance for both C-score<0.5 and C-score<0.3.

**Supplemental Table S1C**. The table shows deletion and duplication predictions of CONGA using Mota, Saqqaq and Yamnaya genomes down-sampled to various depths from their original coverages of 9.6x, 13.1x and 23.3x, respectively. Here, “T” and “F” refer to correct and incorrect predictions respectively, “Miss” is the number of missed true events, “Recall” (TPR) is the true positive rate, and “FDR” is false discovery rate (1 - “Precision”) for each run. The F-Score is calculated as (2 * Precision * Recall) / (Precision + Recall). We calculated “True”, “False”, “Miss”, “Recall”, “Precision”, FDR and F-Score of down-sampled genomes assuming that our CONGA-based predictions with the original genomes (full data) reflect the ground truth. These predictions, in turn, were made using modern-day CNVs as candidate CNV list. The purpose of the experiment was to evaluate accuracy at lower coverage relative to the full data.

**Supplemental Table S1D**. CONGA’s running time and memory consumption on genomes of various depths of coverage calculated using the down-sampled 23x Yamnaya genome (with coverages between 23x and 0.07x) as well as a comparison of CONGA, GenomeSTRiP, FREEC and CNVnator using a 5x simulated genome.

**Supplemental Table S1E**. Results of CONGA runs on simulated genomes at a range of parameters, performed in order to determine the optimum parameters to be used. We tested multiple parameter combinations of C-Score, minimum read-pair support, mappability and minimum mapping quality (MAPQ) using simulated genomes.

**Supplemental Table S1F**. The table contains the data and scripts used to plot figures involving simulated genomes. The graphs were drawn using “ggplot2”.

**Supplemental Table S2**: Information on the ancient genomes used in this study. “ID_publication” refers to the ID of the genome used in the original publication, and “Sample_ID” refers to the ID used in this study. “Average Date (BP)” is the average date in years before present. “Lab PI” refers to the senior author of the study. “Included in final analysis” shows the genomes included in the list of 50 after removing divergent (outlier) genomes.

### 4. Supplemental Notes

**Supplemental Note S1:**

The vast majority of structural variant studies (>92%) available in the European Variation Archive (EVA) are focused on humans (as of March 2022), while the NCBI dbVar has stopped storing non-human SVs. Nevertheless, the EVA database compiles all types of genetic variation data including CNVs from species ranging from the chimpanzee, gorilla, orangutan, rhesus macaque, dog, cow, horse, pig, mouse, zebrafish, and sorghum [14]. Furthermore, CNVs for other organisms are also available in specific databases such as those for *Arabidopsis* [15] and the *Anopheles* mosquito [16].

**Supplemental Note S2:**

Here we discuss the reasons for CONGA's under-performance in genotyping duplications in down-sampling experiments. CONGA's accuracy was particularly low in trials using the Yamnaya genome. This was despite high performance in duplication estimation with simulated genomes. Studying the results in further detail, we noticed that only 47 (2.8%) of the 1,661 originally called duplication events in the full Yamnaya genome were genotyped using read-depth information (i.e. the C-score), while the rest were genotyped using paired-end read information. In contrast, 35% (1498/4027) and 27% (172/638) of duplication events were genotyped using read-depth information on the full Saqqaq and Mota genomes, respectively.

This difference, in turn, can explain CONGA's under-performance at low coverage in the Yamnaya genome: As coverage decreases, the number of paired-end reads supporting a duplication falls rapidly, compromising recall. Meanwhile, the lack of usable read-depth information in the full Yamnaya genome could be related to alignment quality filters applied to the BAM file before data publication. Such filtering likely erased the read-depth signature, leaving only paired-end information available, which is not helpful at low coverages. This scenario is supported by the fact CONGA duplication calls are clearly more successful in the Saqqaq and Mota genomes at low coverage. These two were retrieved as original FASTQ files instead of processed BAM files. The results underscore the need for publishing raw FASTQ files rather than BAM files to allow healthy reuse of the data.

**Supplemental Note S3:**

Here we describe how we selected the divergent, or outlier genome set. We genotyped n=71 real ancient genomes from 10 different laboratories, using all 10,002 human-derived deletions identified in the 1000 Genomes Project African dataset. The presence of technical effects is suggested by a marginally significant laboratory-of-origin effect on deletion frequencies per genome in this set (Kruskal-Wallis test p=0.08). We therefore created heatmaps, multidimensional scaling plots summarizing Euclidean distance matrices, hierarchical clustering trees summarizing Manhattan distance matrices, and principal component analysis plots (PCA) summarizing deletion frequencies (the latter after removing any NAs). We intentionally used a variety of methods in order to capture different potential sources of variability. The results suggested outlier behaviour across n=11 genomes, some that we observed to have relatively low coverage (e.g. ne4, ko2, MA1, DA379), were produced with the capture protocol (Bon002), or originated from the same laboratory despite having variable population origins (e.g. R1, R2, R3, R4, R7, R9; which include Mesolithic and Neolithic genomes ) (also see Supplemental Table S2). Removing these 11 genomes and repeating the analysis with the remaining 60, we again found a significant laboratory-of-origin effect on deletion frequencies per genome (Kruskal-Wallis test p=0.02). The PCA, MDS and hierarchical clustering analyses on the 60 genomes again suggested the presence of a number of divergent cases, mainly from the same laboratories (Iceman, RISE493, RISE495, RISE496, RISE505, RISE511, SI.38, SI.41, SI.45, SI.53). The genome Chan (a European Mesolithic individual) also showed divergent behaviour in the PCA, but this genome has been previously identified as being highly homozygous [17], and it was in fact close to other European Mesolithic individual genomes in its deletion profile, suggesting that the outlier behaviour may be related to its excess homozygosity (indeed this genome had the highest number of homozygous deletions among the 50). We therefore did not remove Chan from the set. We recommend similar quality controls on genotyped deletion datasets created from heterogeneous ancient genomes.
